# Release of human TFIIB from actively transcribing complexes is triggered upon synthesis of 7 nt and 9 nt RNAs

**DOI:** 10.1101/2019.12.19.882902

**Authors:** Elina Ly, Abigail E. Powell, James A. Goodrich, Jennifer F. Kugel

## Abstract

RNA polymerase II (Pol II) and its general transcription factors assemble on the promoters of mRNA genes to form large macromolecular complexes that initiate transcription in a regulated manner. During early transcription these complexes undergo dynamic rearrangement and disassembly as Pol II moves away from the start site of transcription and transitions into elongation. One step in disassembly is the release of the general transcription factor TFIIB, although the mechanism of release and its relationship to the activity of transcribing Pol II is not understood. We developed a single molecule fluorescence transcription system to investigate TFIIB release in vitro. Leveraging our ability to distinguish active from inactive complexes, we found that nearly all transcriptionally active complexes release TFIIB during early transcription. Release is not dependent on the contacts TFIIB makes with its recognition element in promoter DNA. We identified two different points in early transcription at which release is triggered, reflecting heterogeneity across the population of actively transcribing complexes. TFIIB releases after both trigger points with similar kinetics, suggesting the rate of release is independent of the molecular transformations that prompt release. Together our data support the model that TFIIB release is important to maintain the transcriptional activity of Pol II as initiating complexes transition into elongation complexes.

## Introduction

Transcription by RNA polymerase II (Pol II) is a highly regulated process that involves interplay between Pol II, general transcription factors (GTFs), co-regulatory complexes, and gene-specific transcriptional activators and repressors [1–3]. The general transcription factors required to transcribe most protein-coding genes – TFIIA, TFIIB, TFIID, TFIIE, TFIIF, and TFIIH – assemble into preinitiation complexes (PIC) at the promoters of genes with Pol II and co-regulators such as Mediator [4–6]. GTFs play key roles in the formation of PICs, including binding core promoter elements in the DNA, recruitment and correct positioning of Pol II over the transcription start site, and stabilization of Pol II and other factors within PICs [1,7]. Biochemical, structural, and cellular studies have explored the mechanistic roles that each GTF embodies, however, much remains to be learned. This includes developing a better understanding of how GTFs release from complexes as they disassemble and Pol II moves away from the start site of transcription and transitions into elongation.

Toward this goal, we investigated the release of TFIIB from early transcribing complexes. TFIIB is a 33 kDa protein composed of several distinct structural domains that uniquely participate in different aspects of transcription. These include an N-terminal B ribbon region, a B reader domain, a B linker domain, and two cyclin like repeat regions comprising the B core [8,9]. The B core regions directly interact with the TBP subunit of TFIID as well as the upstream core promoter element known as the TFIIB recognition element (BRE), which subsequently positions TFIIB on the DNA in the proper orientation for interactions with Pol II [10]. The B reader domain of TFIIB (composed of the B reader loop, B reader helix, and B reader strand regions) interacts directly with Pol II in a manner that could impact the release of TFIIB during early transcription. A crystal structure of TFIIB bound to initially transcribing Pol II shows the B reader loop positioned in the RNA exit channel leading into the active site of Pol II [11]. A clash between the nascent RNA strand and the B reader loop is predicted to occur once the RNA reaches 7 nt in length, suggesting that TFIIB must undergo a conformational change or release its interactions with Pol II to accommodate the growing transcript [11]. Consistent with this, in vitro transcription assays have shown human TFIIB releases from the promoter DNA during transcription [12–15]; however, it could not be determined whether TFIIB specifically released from transcriptionally active complexes or dissociated from the bulk of inactive complexes in an unregulated manner. Indeed, extensive heterogeneity exists in reconstituted Pol II transcription complexes, and active complexes represent less than 20% of PICs [16–20]. Distinguishing whether TFIIB releases from active complexes or all complexes would reveal whether TFIIB release is an important mechanistic step during early transcription to maintain the activity of Pol II.

Here we determined the relationship between TFIIB release and active transcription complexes. We utilized a single molecule transcription system that can uniquely identify active transcription complexes within a heterogeneous mix of active and inactive complexes [20]. We found that the vast majority of active transcription complexes released TFIIB in an NTP-dependent manner. This correlation and the rate of TFIIB release were not dependent on the interaction between TFIIB and the BRE, the core promoter element to which TFIIB binds. Furthermore, we identified two distinct points in early transcription at which release of TFIIB is triggered: during addition of the seventh and ninth nucleotides to the growing RNA. The rates of TFIIB release after these trigger points are similar. Our data support a model that specific structural changes in transcribing complexes trigger TFIIB to release, then the protein dissociates over time as Pol II continues to transcribe. We propose that release of TFIIB from transcribing complexes is an important switch to maintain the activity of Pol II as it transitions into processive elongation.

## Results and Discussion

### Fluorescently labeled human TFIIB assembles into PICs and supports active transcription

To visualize TFIIB via single molecule microscopy, we created a TFIIB construct containing a C-terminal SNAP tag to allow site-specific labeling with a fluorescent dye [21]. After expression and purification, the TFIIB-SNAP protein was fluorescently labeled with an AlexaFluor488 SNAP dye substrate (AF488-TFIIB). The activity of AF488-TFIIB was compared to untagged TFIIB using a highly purified reconstituted human transcription system. Reactions contained Pol II, TBP, TFIIB, TFIIF, and a core promoter contained on a negatively supercoiled plasmid upstream of a G-less cassette that allowed an RNA product of defined length to be transcribed in the presence of ATP, CTP, and UTP. As shown in Figure 1A, the fluorescent AF488-TFIIB had activity similar to that of the untagged protein, and the system was entirely dependent upon TFIIB.

**Figure 1.**
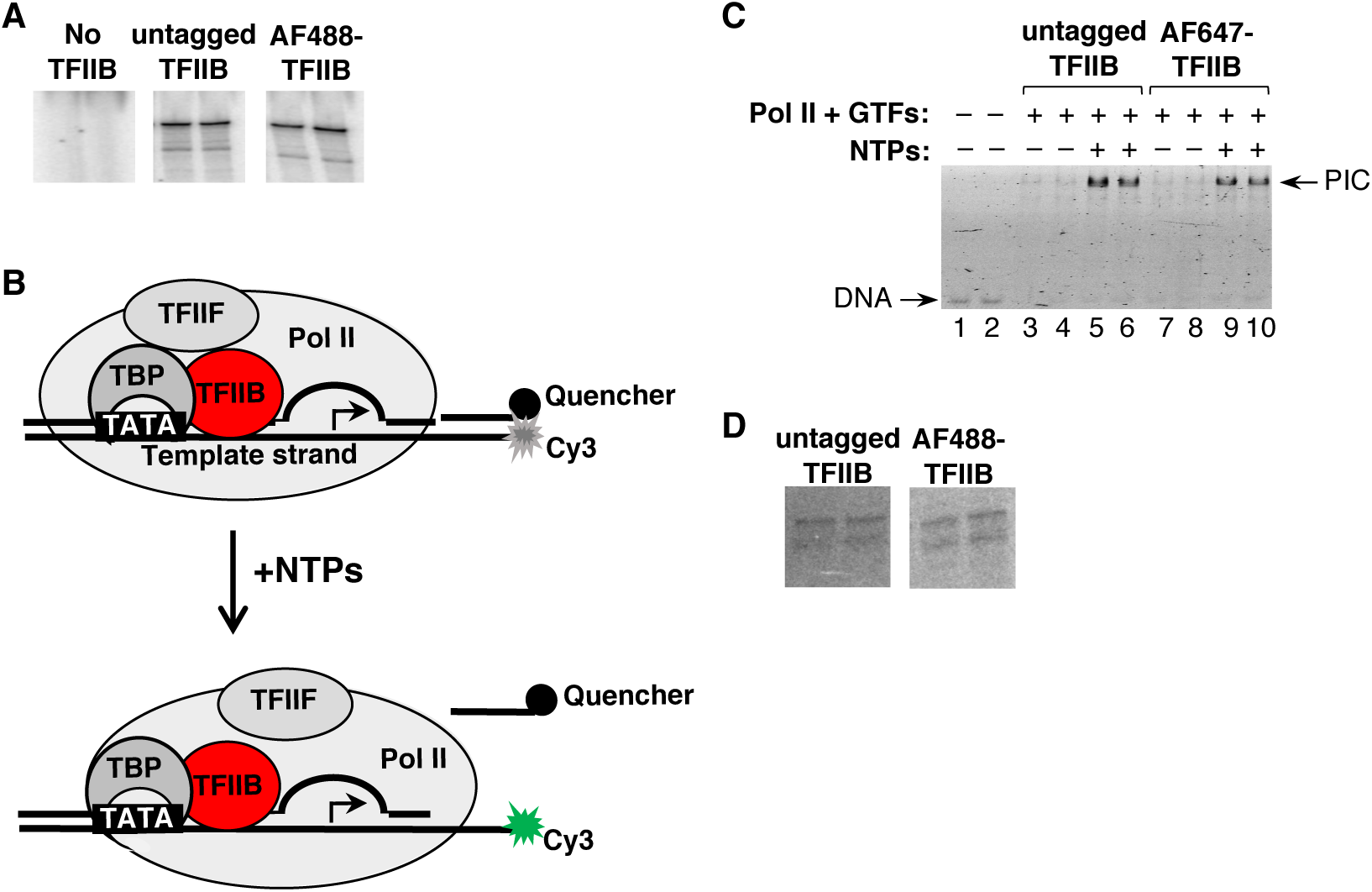
Fluorescent TFIIB-SNAP is active in Pol II transcription assays using a highly purified reconstituted human system. **A.** Untagged TFIIB and AF488-TFIIB show similar activity in ensemble in vitro transcription reactions, and TFIIB is required. Shown are 390 nt G-less transcripts. Each condition was performed in replicate, and the data are from different regions of the same gel at the same exposure. **B.** Schematic of the fluorescence-based transcription assay. PICs are assembled on a DNA construct containing a Cy3 fluorescent dye on the 5’ end of the template strand that is quenched by a Black Hole Quencher 2 molecule attached to a short oligo. After the addition of NTPs, the quencher oligo is displaced by transcribing Pol II, resulting in the appearance of Cy3 fluorescence. Fluorescent AF647-TFIIB is shown in red. **C.** AF647-TFIIB supports PIC formation and active transcription in the quencher-release assay. Transcription complexes were resolved using an EMSA, and the gel was scanned for Cy3 emission. Reactions contained the quenched DNA construct only (lanes 1-2) or PICs formed with either TFIIB (lanes 3-6) or AF647-TFIIB (7-10). NTPs were added to reactions resolved in lanes 5-6 and 9-10. **D.** Similar levels of transcripts are produced from the three piece DNA template with untagged TFIIB or AF488-TFIIB. Shown are RNA transcripts from replicate reactions, and data are from different regions of the same gel at the same exposure.

We next included dye-labeled TFIIB in a fluorescence-based in vitro transcription assay that we previously developed [20]. For this assay, we labeled TFIIB-SNAP with AlexaFluor647 (AF647-TFIIB), which emits in the red channel. As diagrammed in Figure 1B, this assay takes advantage of a three piece DNA construct containing a pre-melted bubble surrounding the transcription start site [22,23]. The template DNA strand has a Cy3 dye on its 5’ end. The non-template DNA strand is composed of two separate oligonucleotides: 1) an upstream oligo that when annealed to the template strand creates the bubble from −9 to +3, and 2) a downstream oligo that contains a BHQ2 quencher molecule on its 3’ end that quenches emission from the Cy3 dye. After addition of NTPs, Pol II transcribes the DNA and displaces the short non-template strand oligo, thereby revealing Cy3 emission.

We coupled this system with electrophoretic mobility shift assays (EMSAs) to evaluate the assembly and activity of PICs containing untagged TFIIB or AF647-TFIIB. PICs were assembled on the three piece DNA construct, then half the reactions received NTPs lacking GTP. The first C residue in the template strand is at +35, hence in the absence of GTP, Pol II pauses at +35 and remains bound to the promoter. All reactions were run in a native polyacrylamide gel (Figure 1C) that was scanned for Cy3 emission, the fluorophore on the DNA. The signal was very faint for unbound DNA (lanes 1-2) and for bound DNA without NTPs (lanes 3-4 and 7-8) due to the non-template strand quencher oligo. The addition of NTPs revealed Cy3 fluorescence (lanes 5-6 and 9-10), reflecting displacement of the quencher oligo due to transcription. This robust NTP-dependent increase in Cy3 emission was observed for PICs formed with either untagged TFIIB or AF647-TFIIB. In addition, similar levels of RNA transcripts were produced from the three piece DNA construct in reactions containing untagged TFIIB or fluorescent TFIIB (Figure 1D).

### Nearly all active transcription complexes release TFIIB

We next investigated the relationship between active transcription and the release of TFIIB by using AF647-TFIIB and the quencher release assay in single molecule experiments. Prior single molecule studies with the quencher-release assay showed that transcriptional activity required the presence of all three GTFs and Pol II, and that ∼10-20% of complexes were active [20]. As diagrammed in Figure 2A, the three piece DNA construct was immobilized on the surface of a streptavidin-derivatized slide using a 5’ biotin on the long non-template strand oligo. This system can resolve active from inactive transcription complexes by identifying individual PICs that contained AF647-TFIIB and also released the quencher oligo after addition of NTPs. To do so, images of surface immobilized Cy3-DNA (green) and AF647-TFIIB (red) molecules before and after addition of NTPs were collected using a TIRF (total internal reflection fluorescence) microscope. Co-localization of red spots on the surface prior to NTPs (i.e. PICs containing AF647-TFIIB) with green spots on the surface after NTPs (i.e. quencher oligo was released) allowed us to identify individual transcription complexes that were active. After identifying active complexes, we inquired about the fate of TFIIB.

**Figure 2.**
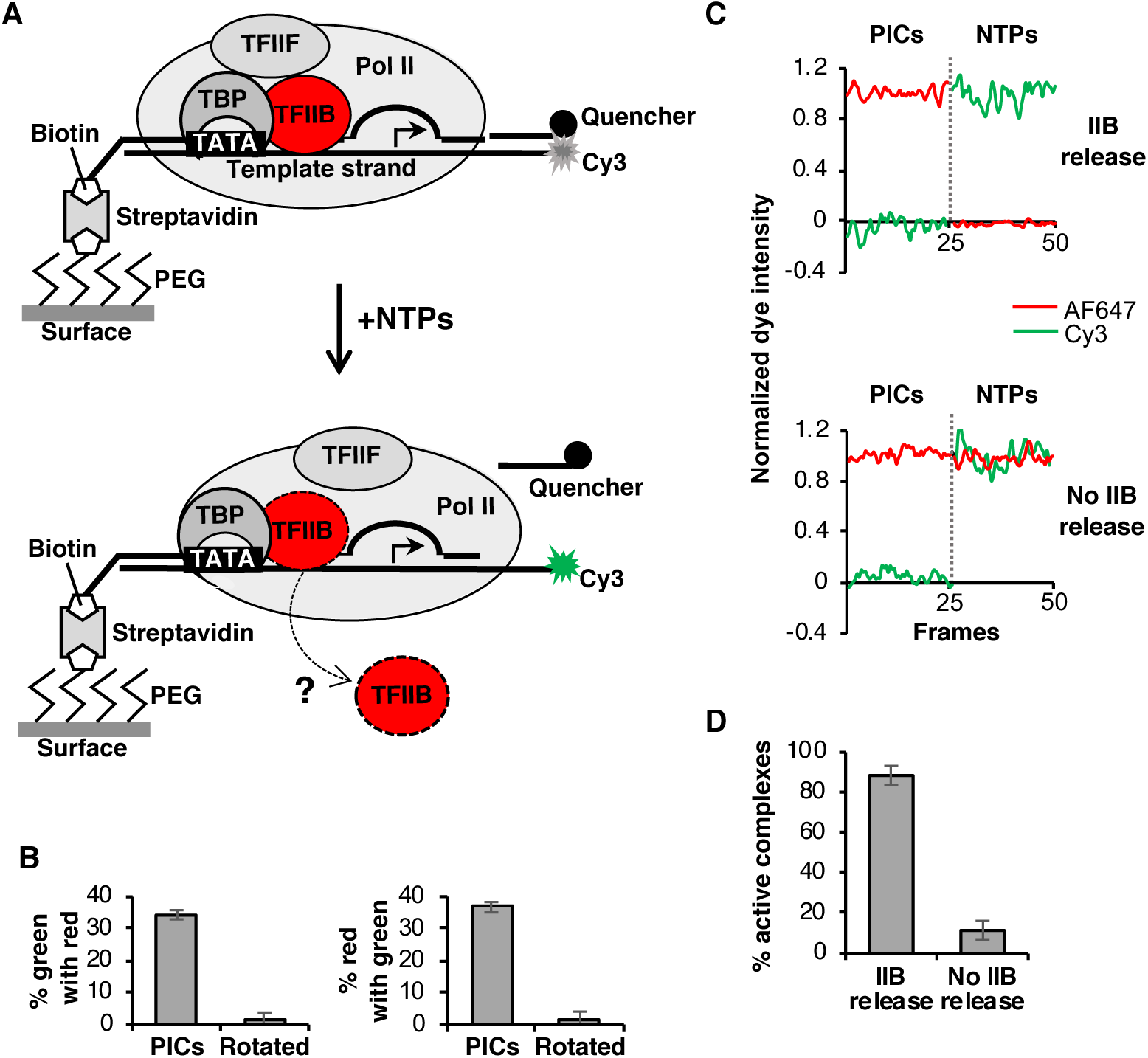
TFIIB releases from nearly all transcriptionally active complexes. **A.** Schematic of the single molecule transcription assay. PICs are immobilized on slides through a biotin on the long non-template strand oligo that binds a streptavidin-derivatized slide surface. The quencher oligo is displaced by transcribing Pol II, resulting in the appearance of Cy3 fluorescence. The fate of AF647-TFIIB in the active complexes can be determined from emission in the red channel. **B.** Single molecule co-localization of AF647-TFIIB emission and Cy3-DNA emission is highly specific. Plotted are AF647 (red) and Cy3 (green) co-localized spots in PICs and after one image was rotated 90°. Each bar is the average of 4 regions and the errors are the standard deviations. **C.** Single molecule intensity traces of Cy3 and AF647 emission from co-localized green/red spot pairs were used to determine the fate of AF647-TFIIB in active transcription complexes after the addition of NTPs. Shown are representative traces from active complexes that either released TFIIB (top) or retained TFIIB (bottom). To create the plots, 25 frames of PIC emission were merged with 25 frames of emission after NTPs. **D.** The majority of transcriptionally active complexes release TFIIB after addition of NTPs. Data are from 248 co-localized spot pairs representing active complexes, plotted is the average of 4 regions and the errors are the standard deviations.

As a control to test the robustness of the red and green co-localization in our system, we first assembled PICs containing AF647-TFIIB on promoter DNA that lacked the quencher oligo so Cy3 emission could be observed in the absence of NTPs. Images of red and green emission were obtained with a TIRF microscope and single PICs were identified using co-localization. We found that 34% of the Cy3-DNA spots (green) co-localized with AF647-TFIIB spots (red), and 36% of the AF647-TFIIB spots co-localized with Cy3-DNA spots (Figure 2B). Importantly, when Cy3 emission images were rotated 90° only 1-2% co-localization of green and red spot pairs was observed, demonstrating that random co-localization in this system is negligible.

Using this single molecule system, we asked whether there was a relationship between the activity of a transcription complex and whether TFIIB released from the DNA after addition of NTPs. PICs containing AF647-TFIIB were immobilized on the surface of a slide and Cy3 (green) and AF647 (red) emission movies were collected with a TIRF microscope. NTPs were added for 2.5 min and a second set of Cy3 and AF647 emission movies were collected over the same regions. We identified the individual PICs that contained AF647-TFIIB prior to addition of NTPs (i.e. red spots on the slide) and were also transcriptionally active upon NTP addition (i.e. green spots after addition of NTPs that co-localized with red spots prior to NTPs). We then determined whether TFIIB released from each active PIC by evaluating whether AF647 emission was lost after the addition of NTPs. AF647 photobleaching was not a concern because slide regions were only imaged for 12.5 s (25 frames with a 0.5 s frame rate) before and after NTP addition.

Figure 2C shows emission data from two different active complexes. On the top, a loss of AF647 intensity (red) was observed after the addition of NTPs, indicating that AF647-TFIIB released from this complex during transcription. The NTP-dependent increase in Cy3 signal (green) showed the complex was transcriptionally active. For comparison, the data on the bottom of Figure 2C were obtained from a complex that was transcriptionally active but did not release AF647-TFIIB during transcription. Here, the Cy3 emission increased with NTP addition, however, the AF647 emission did not change. This analysis was performed for 248 transcriptionally active complexes. As shown in Figure 2D, 88.5% of active complexes released AF647-TFIIB, while only 11.5% retained AF647-TFIIB. To estimate the level of TFIIB release from transcriptionally inactive complexes, we monitored spots of AF647 intensity at which a green dye did not appear upon addition of NTPs. There was not an NTP-dependent increase in AF647-TFIIB released from these transcriptionally inactive spots.

Our data uniquely show that the NTP-dependent release of TFIIB is tightly correlated with transcriptional activity. By making this connection, our single molecule data support the model that release of TFIIB is an important structural re-arrangement that helps maintain Pol II activity as elongation complexes form. Moreover, it is possible that failure to undergo the proper structural transformations that trigger TFIIB release contributes to the high fraction of inactive complexes. Although the vast majority of active complexes released TFIIB, we observed a small fraction of active complexes (∼11%) that retained TFIIB. Interestingly, studies of *E. coli* transcription have shown that the σ^70^ subunit of the polymerase predominantly releases from active transcription complexes; however, it is retained about 30% of the time and might play a regulatory role in transcriptional elongation [24]. There is structural homology between the B reader loop and the 3.2 loop region of σ^70^ [25]. Therefore, it is plausible that retention of TFIIB in a small population of active complexes is a part of an unrecognized regulatory mechanism.

### The rate of TFIIB release from active complexes suggests a trigger mechanism that is not impacted by the BRE

TFIIB can contact the DNA at an upstream core promoter element known as the BRE, which helps position TFIIB in the proper orientation for interaction with Pol II [10]. Accordingly, we asked whether the BRE sequence might influence the relationship between TFIIB release and active transcription. The BRE sequence in our DNA construct is from the adenovirus major late promoter (AdMLP) and differs from consensus at two positions. We made two additional DNA constructs in which the AdMLP BRE was mutated to a perfect consensus sequence (strong BRE) or a completely non-consensus sequence (No BRE). Single molecule transcription assays were performed with the three DNAs to determine the percentage of active complexes that released TFIIB. We found no significant difference between the three promoters (Figure 3A). Hence, the BRE does not influence the tight correlation between active transcription and TFIIB release.

**Figure 3.**
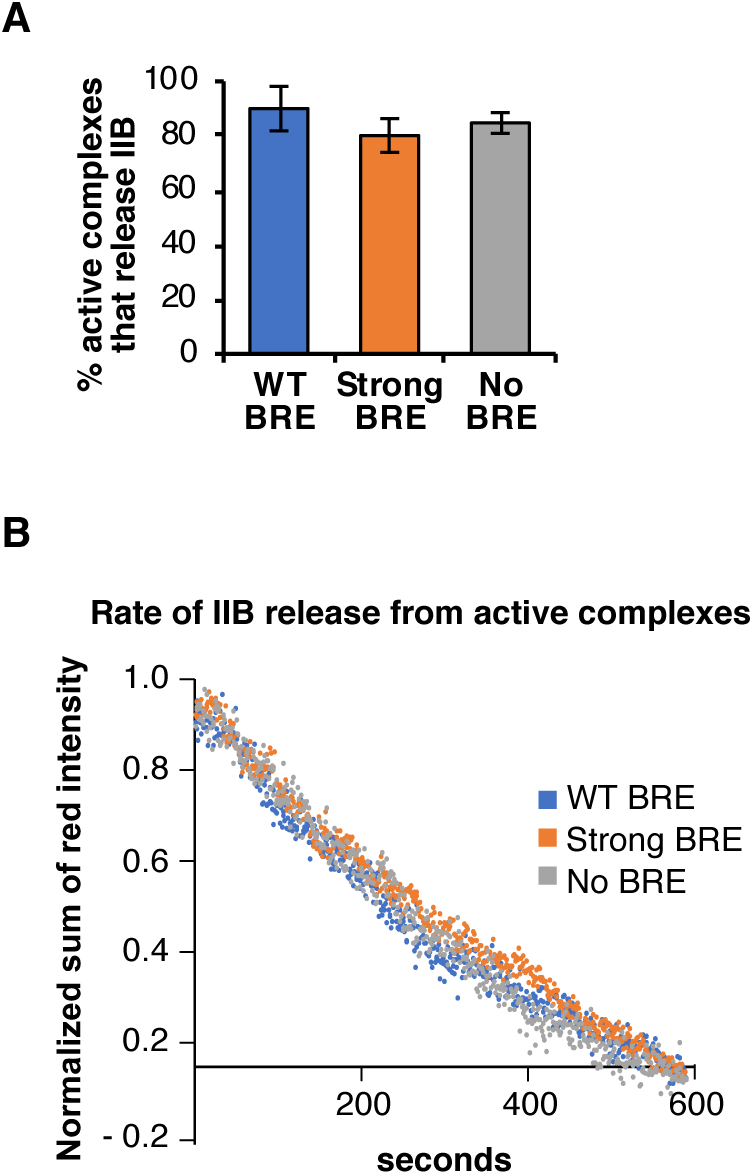
The extent and rate of TFIIB release are not controlled by the BRE. **A.** The percentage of active complexes that released TFIIB was similar with a wild type BRE, a strong BRE, or No BRE; there were 209, 441, and 300 active complexes included in each analysis, respectively. Plotted are the averages of 4 regions and the errors are the standard deviations. **B.** The rate at which TFIIB releases from active complexes is independent of the BRE. For each promoter, the fluorescent intensity of AF647 at each time point from two replicate experiments was summed, then normalized to 1.0 to allow direct comparison of the rate of TFIIB release between templates. There were 91, 62, and 81 active complexes included in the WT, strong, and no BRE analyses, respectively.

These data suggest that whatever role the BRE plays in positioning TFIIB in PICs, it does so in a manner that does not directly impact the activity of transcribing Pol II. In other words, the protein-protein or protein-nucleic acid contacts that must be rearranged to allow TFIIB to release are not controlled by the BRE. The BRE does, however, play a positive role in transcription. Biochemical experiments have shown the BRE enhances transcription levels [10]. Consistent with this, in our single molecule experiments the number of active complexes was ∼30% higher on the strong BRE compared to the No BRE template, suggesting the BRE facilitates the formation of active complexes, but not the release of TFIIB during transcription.

We next asked whether the BRE impacts the rate at which TFIIB releases from complexes during early transcription. We performed single molecule transcription assays on either the wild-type promoter (WT), the strong BRE promoter, or the No BRE promoter. After flowing NTPs over the slide to initiate transcription, AF647 emission was recorded for 10 min to monitor AF647-TFIIB release in real time. Then a short Cy3 emission movie was recorded to identify the DNAs that were transcribed, enabling us to mark the positions of active complexes on the slide surface. We analyzed the data to identify the active transcription complexes that released AF647-TFIIB after NTPs were added. For these complexes, the fluorescence intensity values at each frame of the 10 min movies were summed and plotted versus time to generate a kinetic view of AF647-TFIIB release (Figure 3B). We observed little difference between the three promoters, showing that the BRE does not significantly impact the rate of TFIIB release.

Finding that the BRE also did not impact the rate of release underscores the conclusion that the molecular interactions controlled by the BRE do not regulate TFIIB release. It is most likely that TFIIB makes multifaceted interactions with PIC components, therefore a network of structural changes is required to enable release. Consistent with this, we found that the kinetic data in Figure 3B did not fit well with either a single or double exponential equation, suggesting a multi-step mechanism of release. The half-time for release was ∼300 sec, which is slower than the rate of transcription we previously measured on this promoter using ensemble transcription experiments [26]. This suggests a model in which the release of TFIIB is triggered by an event(s) during early RNA synthesis, then once triggered TFIIB dissociates from complexes at a defined rate as Pol II continues to rapidly transcribe. Interestingly, archaeal transcription factor B (TFB), the homolog of TFIIB, exhibits a multi-step mechanism of release in which the TFB reader domain undergoes structural transformations within complexes between registers +6 and +10, then TFB release is detected at +15 and beyond [27]. It is possible an analogous multi-step mechanism occurs with human TFIIB. Assays that detect conformational changes within active transcription complexes will be required to decipher the structural changes that trigger release of TFIIB.

### TFIIB release is triggered during synthesis of 7 nt and 9 nt RNAs

We next asked whether TFIIB release is triggered at a distinct point or heterogeneously during early transcription. To do so we forced Pol II to pause at specific positions in the early transcribed region using a series of DNAs that each contained a double cytosine mutation in the transcribed region of the template strand. For example, Figure 4A shows the sequence of the template strand of the DNA with cytosines at positions +17 and +18. When ATP, CTP, and UTP were flowed onto the surface of a slide containing PICs assembled on this DNA, Pol II paused after synthesis of a 16 nt RNA due to the lack of GTP. By subsequently flowing in a complete set of nucleotides, Pol II was freed from the pause, elongated the 16 nt RNA to full-length RNA, and displaced the quencher oligo. Pause and release experiments such as these have been widely used to dissect mechanistic steps in Pol II transcription. Using this approach, transcription was paused at eight different positions during early transcription. For each DNA template we sequentially collected three sets of Cy3 and AF647 emission movies from the same regions of a slide in the following order: 1) PICs on the surface prior to addition of NTPs, 2) paused complexes after a 2.5 min incubation with ATP, UTP, and CTP, and 3) complexes after a 5 min incubation with the full set of NTPs. The lasers were only turned on during the short 12.5 s imaging window after each step to remove photobleaching from consideration.

**Figure 4.**
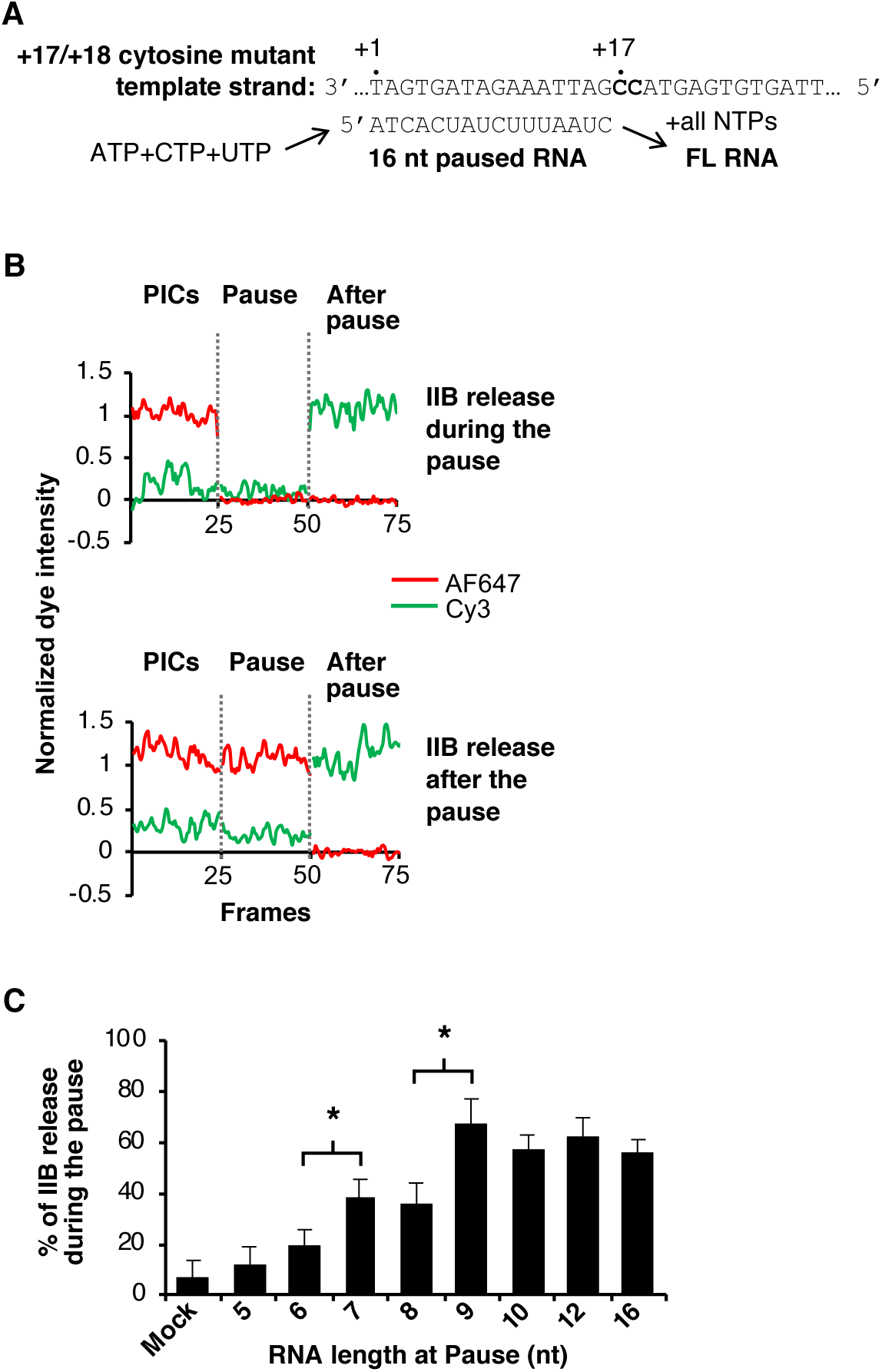
TFIIB release occurs at two distinct points during early transcription. **A.** Cytosine mutant DNA constructs contained two sequential cytosines at various DNA positions on the template strand oligo. The +17/+18 cytosine mutant DNA is shown. On this template, the addition of ATP, CTP, and UTP allows formation of a stable paused complex containing a 16 nt RNA that can be elongated after addition of all NTPs. **B.** Shown are representative intensity traces of Cy3 and AF647 emission for single complexes assembled on the cytosine mutant DNA that allowed Pol II to pause after synthesis of a 7 nt RNA. The data on top revealed that AF647-TFIIB released during the pause, whereas the data on the bottom revealed that AF647-TFIIB released after Pol II was freed from the pause. To create the plots, 25 frames of emission data from each stage of the reaction were merged. **C.** TFIIB release is triggered after synthesis of a 7nt RNA and a 9 nt RNA. The percentage of transcriptionally active complexes that released AF647-TFIIB during the pause was plotted. Data are the average of 8 regions and the errors are the standard deviations. The asterisks indicate where neighboring bars are statistically different (p < 0.01, unpaired two-tailed t-test). The following numbers of active complexes were analyzed, listed according to the RNA length at the pause: 5 nt, 230; 6 nt, 351; 7 nt, 1160; 8 nt, 1482; 9 nt, 655; 10 nt, 127; 12 nt, 258; 16 nt, 401.

Using these emission movies, we first identified spot pairs that showed active transcription and TFIIB release. We then determined whether AF647-TFIIB released during the pause or after Pol II was freed from the pause. Examples of each outcome are shown in the emission traces for two spot pairs (Figure 4B) from complexes assembled on the +8/+9 cytosine mutant template (7 nt paused RNA). In the upper panel AF647-TFIIB released during the pause (i.e. loss of AF647 emission after frame 25 and prior to frame 50). In the lower panel AF647-TFIIB released after Pol II was freed from the pause (i.e. loss of AF647 emission after frame 50). For each of the eight DNA templates that paused Pol II at a different point during early transcription, we identified the active complexes that showed AF647-TFIIB release, then calculated the percentage that released during the pause (Figure 4C). The data show that release of AF647-TFIIB from active transcription complexes was triggered predominantly at two points during early transcription. Specifically, significant jumps in the percent of active complexes that released AF647-TFIIB occurred with synthesis of a 7 nt RNA and a 9 nt RNA. Prior to synthesis of a 7 nt RNA, a low constant level of release was observed, and after synthesis of a 9 nt RNA, the level of release plateaued.

Our data show there are two prominent points at which TFIIB release is triggered during early transcription. This reflects heterogeneity even within the population of active complexes, which would not have been possible to resolve using ensemble experiments. We favor a model in which TFIIB release is triggered by distinct conformational changes within transcribing complexes that occur with synthesis of 7 nt and 9 nt RNAs that serve to destabilize TFIIB contacts with Pol II, and perhaps other GTFs. The first of these conformational changes likely involves removing the TFIIB reader loop from the RNA exit channel of Pol II, as predicted by structural data [11]. Ensemble biochemical experiments have also suggested that destabilization of TFIIB within complexes occurs upon synthesis of a 7 nt RNA [15]. It is less clear what conformational transformations coincide with synthesis of a 9 nt RNA. In addition to breaking contacts between TFIIB and Pol II, these transformations could involve TFIIF, since data show that interactions with TFIIF are critical for holding TFIIB within complexes during initiation [15]. Ensemble biochemical experiments found TFIIB release occurring between synthesis of 10–13 nt RNAs [15]. The differences with our findings could be attributable to the single molecule assays detecting release exclusively from active complexes.

### TFIIB release dissociates from the two populations of complexes with a similar rate

To potentially identify unique characteristics of TFIIB release after the two trigger points, we measured the subsequent rates of release. To do so we paused transcription after synthesis of a 6 nt RNA or an 8 nt RNA, just prior to the trigger points for release, and recorded short Cy3 and AF647 emission movies to mark the positions of complexes on the surface. The complete set of nucleotides was then flowed in to free the paused Pol II, and AF647 emission was recorded for 5 min to monitor AF647-TFIIB release in real time. Lastly, a short Cy3 emission movie was recorded to identify the DNAs that were transcribed. We analyzed the data to identify active transcription complexes that released AF647-TFIIB after Pol II was freed from the pause. For these complexes, the fluorescence intensity values at each frame of the 5 min movies were summed to generate a composite kinetic view of AF647-TFIIB release. The summed AF647 data points were plotted and fit to a single exponential (Figure 5). The average k_release_ values for AF647-TFIIB after freeing Pol II from the 6 nt pause and the 8 nt pause were not substantially different from one another (0.011 ± 0.003 s^-1^ and 0.013 ± 0.003 s^-1^, respectively); the half-times for release are ∼1 min. As a control, we measured the rate at which AF647-TFIIB dissociated from PICs in the absence of NTPs. The half-time was ∼12 min, which is significantly longer than the half-times for NTP-triggered AF647-TFIIB release during early transcription, and more similar to measurements of PIC decay rates [28]. Therefore TFIIB release is transcription-dependent.

**Figure 5.**
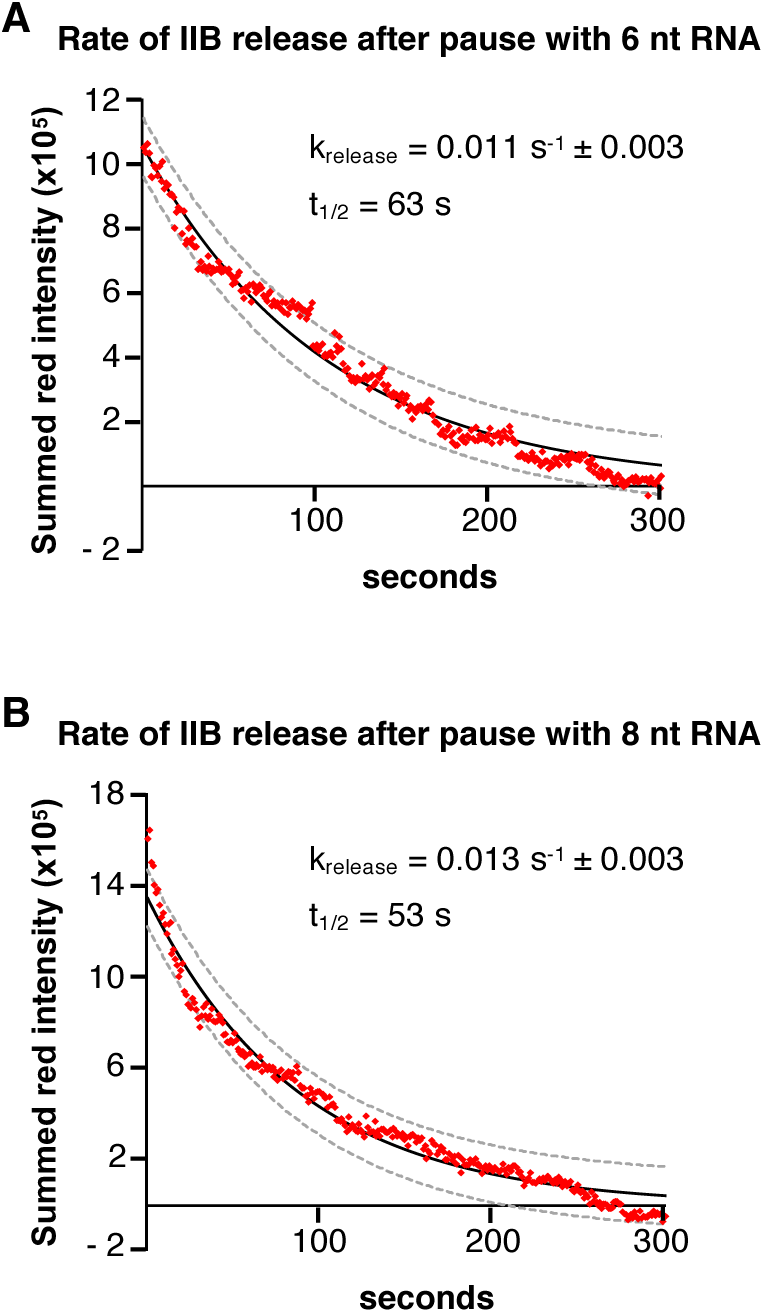
TFIIB releases with a similar rate after the two trigger points in early transcription. Pol II was paused after synthesis of a 6 nt RNA (top plot) or an 8 nt RNA (bottom plot), then the rate of AF647-TFIIB release was measured as Pol II was freed from each pause. Summed AF647 emission (red points) over time was plotted and fit with a single exponential (black lines); the dashed lines represent the 95% confidence intervals of the fits. The average k_release_ values after Pol II was freed from the pauses are shown. The rate constants are the average of two measurements and the errors are the ranges in the data. The t_1/2_ values were calculated from the average k_release_ values. 127 and 95 active complexes were included in each rate measurement for the 6 nt and 8 nt paused RNAs, respectively.

The populations of complexes that release TFIIB after synthesis of a 7 nt or 9 nt RNA are kinetically indistinguishable. Therefore the rate at which TFIIB releases from transcribing complexes is not dependent on the different molecular transformations that trigger its release at the two unique positions. This is consistent with our model that specific structural changes allow, or trigger, TFIIB to release; then TFIIB dissociates with a defined rate as Pol II continues to transcribe the template. This is in contrast to a model in which the polymerase pauses at the trigger points and only continues transcribing after TFIIB release occurs. Interestingly, the rates of TFIIB release we measured after freeing Pol II from paused complexes (Figure 5) are faster than the rate of TFIIB release we measured after the initial addition of NTPs (Figure 3B). This suggests that a kinetically slow step occurs prior to or coincident with synthesis of a 6 nt RNA (the first pause point). The slow step could involve a structural transformation that occurs early after initiation that impacts the positioning of TFIIB in complexes. Our data does not reveal the nature of this transformation, but it could include changes to the network of protein-protein contacts, protein-promoter contacts, or the RNA-DNA hybrid.

Using our single molecule transcription system to resolve active from inactive complexes revealed the tight correlation between transcriptional activity and release of TFIIB. Moreover, we discovered heterogeneity within active complexes that is marked by two different trigger points for TFIIB release during early transcription. We are now positioned to label other general transcription factors to study additional mechanisms by which active PICs disassemble as Pol II transitions into elongation.

## Materials and Methods

### Cloning, expression, purification, and labeling of TFIIB-SNAP

The TFIIB-SNAP construct contained TFIIB followed by an 8-amino acid linker (GSSGGSSG), a SNAP tag, and a His tag. DNAs encoding TFIIB and SNAP were amplified using PCR and inserted into a pET21a+ expression vector using NdeI and BamHI restriction sites. TFIIB-SNAP was expressed in BL21 *E. coli* cells. The cell pellet was lysed by sonication in Buffer A (20 mM Tris pH 7.9, 500 mM NaCl, 10% glycerol, 2 mM DTT, 0.5 mM PMSF, and 1X EDTA-free Protease Inhibitor (Roche)) and centrifuged. The supernatant was loaded on a HisPur Ni-NTA (Thermo Scientific) column pre-equilibrated with Buffer A. The column was washed multiple times with Buffer A followed by Buffer A containing 50 mM imidazole. TFIIB-SNAP was eluted with Buffer A containing 300 mM imidazole. The eluate was dialyzed overnight at 4°C into 10 mM Tris pH 7.9, 50 mM KCl, 1 mM DTT, and 10% glycerol. To fluorescently label the protein, 100 µl of 2 µM TFIIB-SNAP was incubated with 10 µM SNAP-Surface AlexaFluor dye substrate (New England Biolabs) for 2 hr at room temperature in the dark. Excess free dye was removed using Zeba Spin Desalting columns (7K MWCO, 0.5 mL, Thermo Scientific).

### Three piece DNA constructs for fluorescent transcription assays

The DNA contained the AdMLP from −40 to −1 and a downstream sequence optimized for template usage, as previously described [20]. Constructs were generated by annealing three oligos (the BRE is underlined and the start site is in bold): 1) template strand oligo (−40 to +37) containing a 3’ Cy3 fluorophore: 5’CCTGAGGTTAGTGTGAGTAGTGATTAAAGATAGTGA**T**GAGGACGAACGCGCCCCC ACCCCCTTTTATA<u>GCCCCCC</u>TT3’, 2) long non-template strand oligo (−40 to +20): 5’AA<u>GGGGGGC</u>TATAAAAGGGGGTGGGGGCGCGAAGCAGGAG**T**AGACTATCTTTAA TCACTA3’, and 3) short non-template strand oligo (+21 to +37) containing a 3’ BHQ2 quencher: 5’CTCACACTAACCTCAGG3’. The first C residue in the template strand is at +35, hence in the absence of GTP, Pol II pauses at +35. The sequence of the non-template and template strands from −9 to +3 will not anneal in order to create a pre-melted region. For single molecule experiments the long non-template strand oligo contained 5’-biotin-TACGAGGAAT to enable surface immobilization. For the BRE mutants, the AdMLP BRE sequence (GGGGGGC, non-template strand underlined above) was changed to GGGCGCC and AACTCGG for the strong BRE and No BRE, respectively. For pausing experiments, the following DNA positions were individually mutated to CC on the template strand and GG on the non-template strand: +7/+8, +8/+9, +9/+10, +10/+11, +11/+12, +13/+14, +17/+18. All oligos except the short non-template strand oligo were gel purified prior to annealing. In annealing reactions the molar ratio of template oligo:long non-template oligo:short non-template oligo was 2:1:10. The oligos were annealed in a thermocycler as follows: 5 min at 95°C, cool to 60°C at 0.1°C per sec, 1h at 60°C, cool to 45°C at 0.1°C per sec, 1h at 45°C, cool to 4°C at 0.1°C per sec.

### Ensemble transcription assays

Recombinant human TBP, untagged TFIIB, TFIIF, and native human Pol II were prepared as described previously [20]. The ensemble in vitro transcription reactions were assembled as previously described [20]. Briefly, reactions were performed in 10% glycerol, 10 mM Tris (pH 7.9), 50 mM KCl, 1 mM DTT, 0.05 mg/ml BSA, 10 mM HEPES (pH 7.9) and 4 mM MgCl_2_. Reactions contained 1-2 nM DNA, either the three piece construct or plasmid DNA containing the AdMLP fused to a 390 bp G-less cassette [29]. Proteins were added to the reactions at final concentrations of 3.5 nM TBP, 2 nM TFIIF, 2 nM Pol II, and 10 nM untagged TFIIB or fluorescently labeled TFIIB-SNAP. PICs were formed by incubation for 20 min at room temperature, then NTPs were added for 20 min (625 µM ATP, UTP, and 25 µM [^32^P]CTP, 5 Ci per reaction). Reactions were stopped and RNA transcripts were ethanol precipitated as described [16]. The RNA was resolved in 6% (Figure 1A) or 20% (Figure 1D) denaturing gels. To observe the formation of PICs and quencher oligo release (Figure 1C), reactions were run on 4% native gels containing 0.5X TBE and 5% glycerol [20]. Native gels were imaged by scanning with a 532 nm and laser and reading fluorescence emission using a 580 nm emission filter on a Typhoon Imager 9400 (GE biosciences).

### Single molecule transcription assays

The cleaning and assembly of the flow chambers, the surface functionalization, and the preparation of streptavidin, D-glucose, glucose oxidase, catalase, and 100 mM Trolox stock solutions were as previously described [30]. Preinitiation complexes were formed by incubating 0.15-0.25 nM biotinylated three piece DNA with 3.5 nM TBP, 5 nM AF647-TFIIB, 2 nM TFIIF, 2-3 nM Pol II, and 0.5 ng/µl poly(dG:dC) competitor DNA in DB/RM buffer (10 mM Tris pH 7.9, 10 mM HEPES pH 7.9, 50 mM KCl, 4 mM MgCl_2_, 10% glycerol, 1 mM DTT, 0.1 mg/ml BSA) for 20 min at room temperature. PICs were then flowed onto the surface and incubated for 10 min. The surface was washed twice with DB/RM buffer, then Imaging Buffer (1.02 mg/ml glucose oxidase, 0.04 mg/ml catalase, 0.83% D-glucose, and 3.45 mM Trolox in DB/RM buffer) was flowed into slide chambers. For non-pausing transcription assays NTPs (625 µM ATP, CTP, and UTP) diluted in Imaging Buffer were flowed into slide chambers and incubated for 5 min. For pausing assays all NTP solutions were made in Imaging Buffer containing 1.6 U/µl of Murine RNase Inhibitor (New England Biolabs). To pause after synthesis of a 5 nt RNA, PICs were assembled on wild type DNA and 1 mM ApU, 625 µM ATP and 625 µM CTP were added for 2.5 min. To pause transcription at other positions, PICs were assembled on cytosine mutant DNAs and 625 µM ATP, CTP, and UTP were added for 2.5 min. For the mock pause control, a solution lacking NTPs was used. In all cases, the pause was released by incubation with a solution of 625 µM ATP, CTP, UTP, and GTP for 5 min.

### Single molecule data collection and analysis

Flow chambers were imaged using an objective based TIRF microscope (Nikon TE-2000U) equipped with a 1.49 NA immersion objective, a Piezo nanopositioning stage, and two CCD cameras. Samples were excited with 532 nm (green) and 635 nm (red) continuous wave lasers, and emission images were collected by an Evolve Photometric CCD and a Cascade II Photometric CCD, respectively. Green and red emission movies were sequentially recorded using the NIS-elements software at 0.5 sec exposure time for 25 frames, except for the kinetic experiments, which used a 1 sec exposure time over the time frames noted in the Results and Discussion. By using the Piezo nanopositioning stage, green and red emission movies across the same regions were recorded for PICs and after NTP addition. These movies were analyzed with in-house co-localization software to identify green/red spot pairs representing quencher released from PICs with AF647-TFIIB. As a control, co-localization analyses were also performed with the Cy3 image rotated by 90° to ensure that co-localization was not due to random overlap of spots. Fluorescence emission traces for every co-localized spot pair were individually viewed and analyzed. The green intensity traces for each spot pair were used to identify active transcription complexes via an increase in Cy3 emission due to quencher release after addition of NTPs. The red intensity traces for each spot pair were used to determine if and when TFIIB released from Pol II via a decrease in AF647 emission.

## Acknowledgements

This work was supported by the National Science Foundation (MCB-1817442 and DGE1144083 to E.L.) and the National Institutes of Health (T32 GM08759 to E.L. and T32 GM065103 to A.E.P).

## Author contributions

Conceptualization: E.L., A.E.P., J.A.G., J.F.K.; Reagent preparation and single molecule data collection: E.L., A.E.P.; Data analyses: E.L., A.E.P., J.A.G., J.F.K.; Funding acquisition: E.L., J.A.G., J.F.K.; Software: J.A.G.; Supervision: J.A.G., J.F.K.; Writing: E.L., J.A.G., J.F.K.; All authors edited and approved of the final manuscript.

## Abbreviations

Pol II: RNA polymerase II
GTF: general transcription factor
TFII-: transcription factor of Pol II
PIC: preinitiation complex
AF488-TFIIB: TFIIB labeled with AlexaFluor488
AF647-TFIIB: TFIIB labeled with AlexaFluor647
BRE: TFIIB recognition element
EMSA: electrophoretic mobility shift assay
TIRF: total internal reflection fluorescence
AdMLP: adenovirus major late promoter.

